# An Operant-based Touchscreen Morph Discrimination Task Does Not Detect Age-related Mnemonic Similarity Deficits in Rats

**DOI:** 10.64898/2026.04.30.722044

**Authors:** Aleyna Ross, Carly N. Logan, John J. Thompson, Sarah A. Johnson, Cory Watson, Maria Ramirez, Katelyn N. Lubke, Andrew P. Maurer, Sara N. Burke

## Abstract

The Mnemonic Similarity Task (MST) is highly sensitive to age-related cognitive decline in humans and has been adapted for rodents using 3D objects, where aged animals show deficits in discriminating similar lures. To improve translational alignment with human testing and increase automation, we developed a touchscreen-based rat analog using a morphed Object-Cued Spatial Choice (OCSC) task with 2D image stimuli. Young (4-month) and aged (21-month) male and female Fischer 344 × Brown Norway hybrid rats were trained in Bussey-Saksida touchscreen chambers and tested on discrimination performance using image pairs that varied parametrically in feature overlap. We also assessed perirhinal cortical engagement in a subset of animals using *Arc* expression as a readout of activity-related principal cell firing following low-and high-overlap task epochs. Across shaping and procedural training, aged rats required more errors to reach criterion on one stimulus set, but both age groups successfully acquired the task. During morph testing, performance declined systematically as stimulus similarity increased, confirming that the task manipulated discrimination difficulty. However, contrary to expectations, young and aged rats performed similarly across overlap conditions, with no significant age-related impairment. In the *Arc* experiment, discrimination accuracy was again reduced by greater stimulus overlap, but *Arc* expression in perirhinal cortex did not differ reliably by age or overlap condition, although expression was associated with behavioral accuracy and deep layers showed higher ensemble similarity than superficial layers. These findings indicate that, while the touchscreen morph OCSC task is sensitive to stimulus similarity, it does not detect the robust age-related mnemonic discrimination deficits previously observed with 3D object-based rodent MST paradigms, underscoring the importance of considering ethological relevance when designing translational cognitive assays.

## INTRODUCTION

To optimize the health and wellbeing of older adults, early identification of subtle shifts in cognition is critical for offering the best opportunity to delay or even prevent the onset of more debilitating cognitive impairment. One task that is particularly sensitive to detecting early age-related cognitive change (Yassa et al., 2010; Stark et al., 2015; Stark and Stark, 2017; Stark et al., 2019) is the Mnemonic Similarity Task (MST), which quantifies a study participant’s accuracy in discriminating between familiar stimuli and novel lures that are similar to the familiar items (Kirwan and Stark, 2007; Stark et al., 2013). In humans, performance impairments on the MST are associated with dysfunction in hippocampal-cortical circuits (Yassa et al., 2011b; Yassa et al., 2011a; Bennett et al., 2015; Bennett and Stark, 2016; Reagh et al., 2018; Bennett et al., 2019; Stark et al., 2019) that vulnerable in advanced age (for review, see Burke and Barnes, 2024). For example, MST impairments are associated with an elevated BOLD signal in the CA3/dentate gyrus subregions of the hippocampus (Yassa et al., 2011b), lower activation in anterolateral entorhinal cortex (Reagh et al., 2018), and a hyperconnectivity between the anterolateral entorhinal cortex and CA3/dentate gyrus (Adams et al., 2022). Furthermore, loss of perforant path and fornix fibers that connect the hippocampus to cortical and subcortical structures, respectively, is also related to impaired MST performance (Bennett et al., 2015; Bennett and Stark, 2016). Recently, the MST has been adapted for quick web-based assessment (Stark et al., 2023), and when granular analyses are applied that take in account participant response biases, it can predict biomarker levels for Alzheimer’s disease related pathology (Vanderlip et al., 2024).

The promise of the MST as a digital biomarker points to the utility of developing a modified version of this task for use in preclinical animal models to interrogate mechanisms and design more effective interventions. The first study to incorporate target-lure discrimination abilities in animal models used objects with shared features to test novel-familiar object discrimination in a spontaneous novel object recognition task (Ennaceur and Delacour, 1988). Although young rats showed a preference for the novel object, aged rats falsely recognized a novel lure object as familiar, failing to exhibit lure discrimination (Burke et al., 2011). This same study also constructed objects from LEGO® blocks to parametrically vary stimulus overlap and reported that aged bonnet macaques could not discriminate between objects with feature overlap to the same degree as young monkeys (Burke et al., 2011). More recently, a lure discrimination task for rats was implemented to specifically parallel the MST. Comparable to the encoding phase of the human MST, rats were trained to respond to a familiar LEGO object target. Once this easy discrimination problem was learned, during the testing phase, the learned target was presented with one of the novel lures and the animal was rewarded for selecting the target. Young and aged rats were similarly able to select the target over dissimilar lure objects, but aged rats were selectively impaired at discriminating the target from similar lures (Johnson et al., 2017), as observed in humans. The rodent MST has also been tested using odorant stimuli. Within a homologous chemical series (e.g., alcohols or aldehydes), feature overlap between two odorants can be manipulated by varying the number of carbon atoms between the 2 odorants (Yoder et al., 2014). Young and aged rats have similar odor detection thresholds and discrimination accuracy for odorants of different chemical classes or when the carbon lengths of the two odorants varies by 5. When the carbon lengths only differ by 1 or 3 carbons, however, aged rats are significantly less accurate than the young (Yoder et al., 2017), indicating that age-associated mnemonic discrimination deficits are sensory modality invariant.

The rodent MST has allowed the causative relationship between hippocampal-cortical circuitry and performance to be interrogated, offering validity to the human studies that have reported associations between neurobiology and mnemonic discrimination. Specifically, elevating activity in the dorsal CA3/dentate gyrus (Johnson et al., 2019), or disrupting cortical inputs to the hippocampus with a unilateral knife cute of the perforant pathway (Burke et al., 2018a) causes a decline in MST performance in young rats. While the findings from studies using animal models and human participants demonstrate parallels, one striking difference between the rodent and human variants of the MST are the types of stimuli used for testing. The human MST studies employ 2D images of everyday objects, whereas the rodent MST uses 3D objects in arenas or mazes (Johnson et al., 2017). This discrepancy could create a significant gap in translational relevance, as the two formats differ in complexity and stimulus control. Additionally, the LEGO-based MST demands substantial manual involvement from the experimenter, and typically only permits one rat to be tested at a time.

Advances in the ability to use touchscreen operant chambers to test rodents have enabled the automated presentation of 2D image stimuli and rapid response collection, eliminating the need for manual input, reducing variability and potential enhancing reproducibility and rigor (Morton et al., 2006; Bussey et al., 2008). Moreover, touchscreen based cognitive testing of Paired Associated Learning performance has established striking parallels between humans and rodents for impairments related to age (Robbins et al., 1994; Lee et al., 2013; Smith et al., 2021), traumatic brain injury (Newcombe et al., 2016; Smith et al., 2023) and schizophrenia (Nithianantharajah et al., 2015). These studies highlight the potential of touchscreen cognitive testing across species for bridging the translation gap between preclinical and clinical research. Furthermore, the use of 2D image stimuli allows for greater flexibility and better alignment with the stimulus formats used in human MST studies. The goal of the current study was to design a touchscreen-based version of the MST using morph stimuli. The object-cued spatial choice (OCSC) task (Ahn and Lee, 2015) was adapted in an attempt to design a touchscreen based MST. This task was selected because it requires activity in the perirhinal cortex (Ahn and Lee, 2015), a structure that projects directly to the hippocampus (Naber et al., 1999) that is involved in stimulus recognition and discriminating between stimuli that share features (Bartko et al., 2007b, a). In the morphed OCSC task, yyoung and aged rats both showed declining performance when feature overlap between stimuli increased, however, there were no behavioral differences between age groups and therefore this novel task design cannot be considered a new rat analog of the MST. These observations highlight the importance of balancing comparable testing platforms across species with ethological relevance.

## METHODS

### Animals and handling

Young (4-month, n = 11♀/11♂) and aged (21-month, n = 10♀/8♂) male and female Fischer 344 x Brown Norway hybrid rats were used in this study. Animals were single housed in standard Plexiglas cages and kept on a reverse 12-hour light/dark cycle, with all testing occurring during the dark/active phase. For enrichment purposes, each cage was furnished with a retreat box and Nyla bone. During the first week of habituation to the University of Florida (UF) vivarium, all animals were provided with ad libitum food and water. One week after arrival at the facility, rats handled daily for 3-5 days for acclimation to experimenters as well as basic procedures that included transporting rats to the testing room on a cart and picking up rats by the midriff to get accustomed to experimenter interaction and daily weighing. Rats were also placed on food restriction to encourage motivation on the appetitively reinforced task. During restriction, water access remained ad libitum and rats received 15-30 g (∼1.9 KCal/g) moist chow once daily with the amount being adjusted based on the animal’s current weight and their body condition score. The body condition score is assigned based on the presence of palpable fat deposits over the lumbar vertebrae and pelvic bones (Hickman and Swan, 2010; Siriarchavatana et al., 2022). As in our previous publications (Johnson et al., 2017; Smith et al., 2021), the baseline weight was determined to be the weight that corresponded to an optimal body condition score of 3 with additional considerations given to aged males (Logan et al., 2025). For young male and female animals this tended to be the initial weight upon arrival at the facility. Aged male rats arrive over conditioned from prolonged ad libitum feeding (Martin et al., 2010), and typically require >65 grams of weight loss to reach an optimal body condition. Once the rats reached 85% of their baseline/ideal BSC weight, behavioral shaping procedures began. All procedures were in accordance with the NIH Guide for the Care and Use of Laboratory Animals and approved by the Institutional Animal Care and Use Committee at the University of Florida.

### Apparatus

The automated morph object-cued spatial choice (OCSC) task was conducted in a behavioral testing chamber equipped with a touchscreen interface known as Bussey-Saksida Touchscreen Chambers for Rats. The touchscreen operant chamber consisted of an opaque Perspex box mounted in a sound-attenuating cupboard. Bussey-Saksida touchscreen operant chambers were purchased though Lafayette Instrument (Lafayette, IN) and manufactured by Campden Instruments (Loughbrorough, UK). A touch-sensitive screen (15 in x 12 in) responsive to a ‘nose-poke’ was located at one end of the box. After initial shaping, the areas of responsive touchscreen were restricted by covering the screen with a black plastic mask with three windows that framed the presentation zones of the stimuli. The mask also contained a shelf that provided a desk-like platform to encourage rats to stand on their hind paws and view the stimuli presented in the response windows as well as to prevent accidental touches. A feeder connected via clear PVC tubing (Fisherbrand™) to a pump and reservoir containing liquid reward (Ensure® Strawberry Nutrition Shake, Abbott Laboratories) was located at the opposite end of the chamber. In all shaping, training, and testing protocols, session start was signaled by illumination of a house light mounted on the chamber ceiling and the playing of a tone through a speaker mounted on a shelf above the chamber. After the first trial was initiated, the chamber was then kept dark during the session except when the rat made an error, the house illuminated to indicate the incorrect response. Infrared (IR) beams positioned in the feeder and at the front and rear of the chamber register the rat’s position during each session. Control of outputs (including house light, feeder light, stimulus presentation, tones, and reward dispenser) and collection of data from inputs (IR beams and touchscreen responses) was implemented with a custom software interface supplied with the operant chambers (ABET II software, Lafayette Instrument Co.).

### Shaping

Prior to Morph task testing, animals underwent a pre-training phase to acclimate to the touchscreen apparatus and to learn the basic task contingencies through shaping procedures for the Object-cued Spatial Choice (OCSC) task. Shaping to familiarize rats with the components of the touchscreen chambers and the basic trial structure proceeded in six phases: 1) Magazine Training, 2) Any Touch Full Screen and Progressive, 3) Any Touch Mask Only, 4) Must Touch, 5) Must Initiate, and 6) Punish Incorrect. *See* ***Table 1*** *for summary statistics of each shaping phase.* These shaping procedures were standardized across most protocols supplied with the ABET II software package and were adapted here from shaping schedules provided with the Paired Associate Learning (PAL) task (Bussey et al., 2009; Bussey et al., 2015). The initial stage of shaping, Magazine training (**Figure1A**), involved training rats to retrieve the liquid reward. This also ensured that food restriction was sufficient to appetitively motivate the animals. **Figure 1B** shows the average number of days required by age group to reach criterion on magazine training.

**Figure 1.**
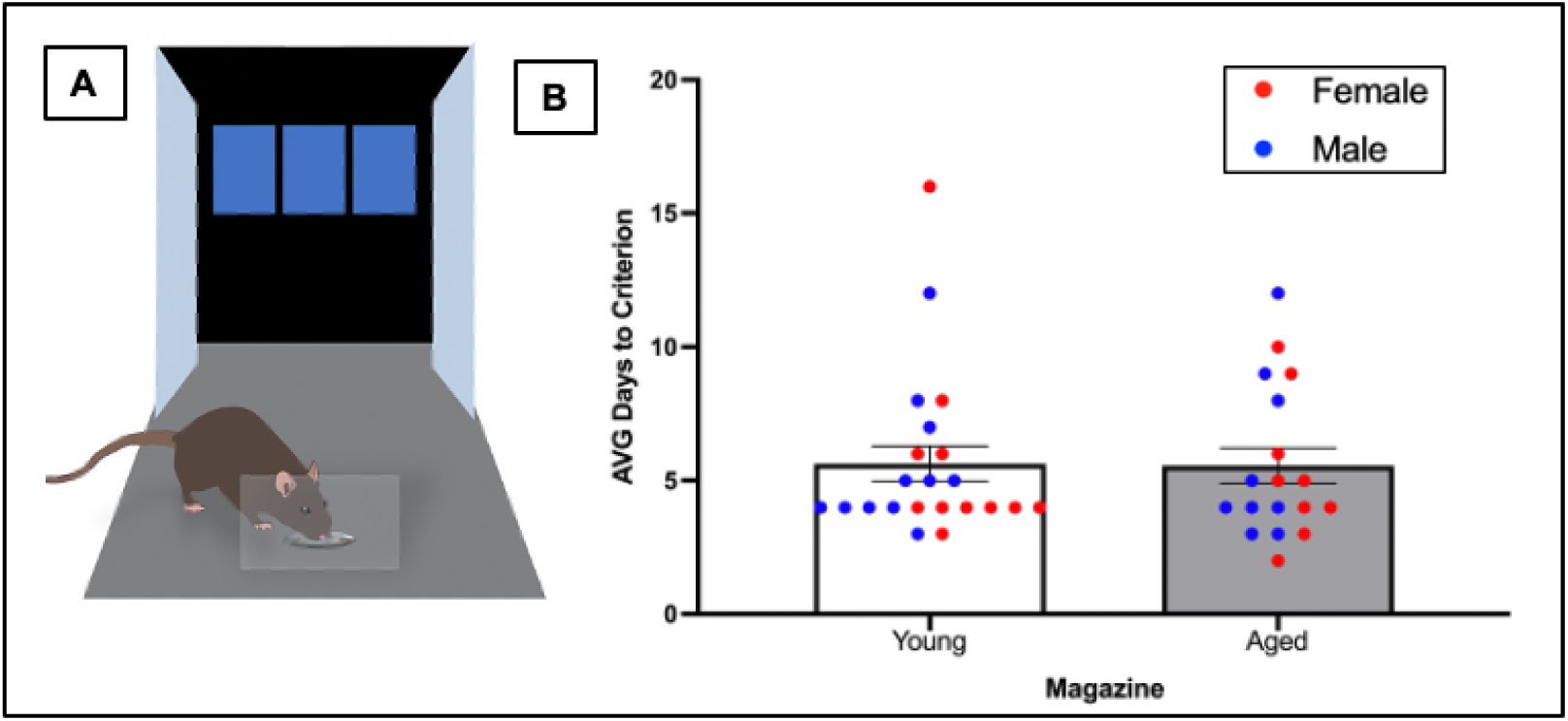
Magazine training in the Object-Cued Spatial Choice Testing task. (A) Representative image of a rat undergoing magazine training in Touchscreen operant chamber. During this phase, rats learned to associate the food magazine with reward delivery. (B) Average number of days required to reach the magazine training criterion in young and aged rats. Data are presented as mean ± SEM with individual data points overlaid and color-coded by sex.

**Table 1.** Summary Statistics of OCSC Shaping Procedures, *p<0.05.

| OCSC Task Shaping Procedures | Sex | Age | Age x Sex |
| --- | --- | --- | --- |
| **Stage 1** |  |  |  |
| Magazine Training | $F(1,36)=0.021, p=0.886$ | $F(1,36)=0.008, p=0.928$ | $F(1,36)=0.002, p=0.967$ |
| **Stage 2** |  |  |  |
| Fullscreen | $F(1,36)=2.603, p=0.115$ | $F(1,36)=5.824, p=0.021 *$ | $F(1,36)=0.430, p=0.516$ |
| Progressive | $F(1,16)=1.822, p=0.196$ | $F(1,16)=1.860, p=0.192$ | $F(1,16)=1.200, p=0.290$ |
| Mask Only | $F(1,34)=0.928, p=0.342$ | $F(1,34)=1.981, p=0.168$ | $F(1,34)=0.092, p=0.764$ |
| **Stage 3** |  |  |  |
| Must Touch | $F(1,36)=3.118, p=0.086$ | $F(1,36)=0.904, p=0.348$ | $F(1,36)=1.094, p=0.303$ |
| Must Initiate | $F(1,36)=0.698, p=0.409$ | $F(1,36)=7.327, p=0.010 *$ | $F(1,36)=0.086, p=0.771$ |
| Punish Incorrect | $F(1,36)=0.046, p=0.832$ | $F(1,36)=2.563, p=0.118$ | $F(1,36)=0.016, p=0.899$ |

During Any Touch shaping, the objective was to encourage rats to interact with the touchscreen by rewarding nose-poking at any location on the screen. This phase of shaping was adapted to include three sub-phases, which in our experience, sped up the rats’ progress. These sub-phases were referred to as Any Touch – Full Screen, Progressive, and Mask Only. During the Full Screen (**Figure2A**, left panel) sub-stage, the mask was not used, and the entire screen was exposed **i**n each trial, the screen was automatically illuminated, and the rat could nose-poke anywhere on the screen to initiate the delivery of a liquid reward. If the rat failed to interact with the screen, no reward was delivered. Once the rat reached a criterion of obtaining 100 rewards in a 45-minute session, the animal proceeded to the next sub-phase of Any Touch shaping (Progressive; **Figure 2A**, middle panel). In this phase of the experiment, the shelf was placed in the touchscreen, covering the bottom half, and the top half of the screen was illuminated to decrease the amount of available touch area and encourage the animal to touch at a higher level. Any nose-poke on the illuminated touchscreen resulted in a liquid reward. Once the rat reached 100 rewards obtained, it proceeded to the next phase of the study (Mask Only; **Figure 2A**, right panel). During Mask Only training, the available space to touch the screen was confined to the three response windows and the rest of the screen was blocked by the mask. The rat was required to touch any position in the center, leftmost, or rightmost window to receive a liquid reward until they were able to retrieve 100 rewards. **Figure2B** shows the mean number of days that young and aged rats spent at each sub-phase of Any Touch shaping to reach criterion.

**Figure 2.**
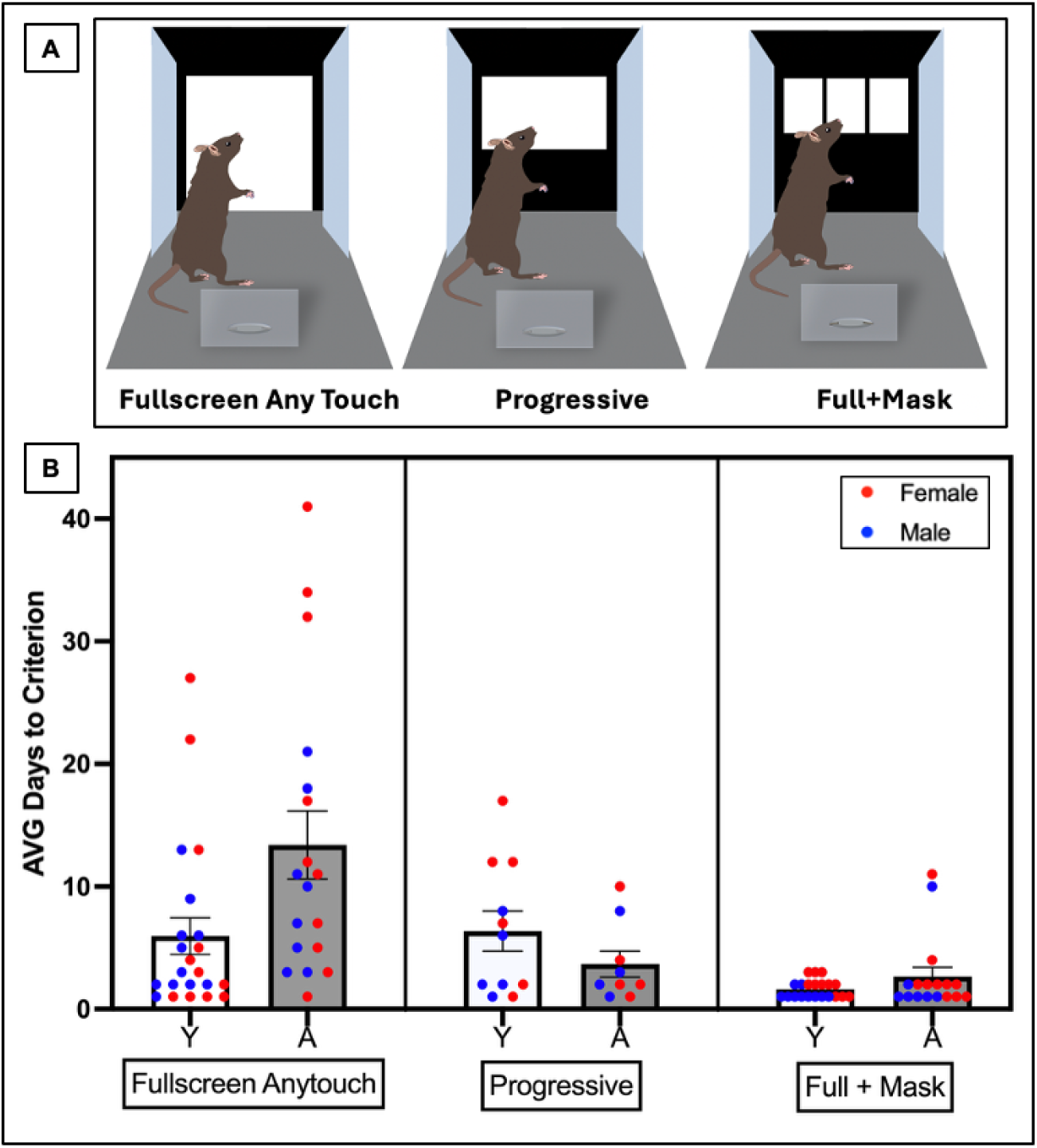
Any Touch Shaping. (A) Representative images illustrating the three shaping phases: Fullscreen Anytouch, Progressive, and Full+Mask. In the Fullscreen Anytouch phase, the entire touchscreen is responsive, allowing the rat to learn the screen-touching response. The Progressive phase reduces the responsive area to the top half of the screen, reinforcing directed nose-poke behavior. The Full+Mask phase introduces a physical mask that reveals only three response windows, refining spatially targeted responding. (B) Average number of days required to reach criterion for each shaping phase in young (Y) and aged (A) rats. Data are presented as mean ± SEM with individual data points overlaid and color-coded by sex.

In the next training phase, “Must Touch”, only one response window of the three within the screen mask was illuminated (**Figure 3A**, left panel). The rat was required to touch the illuminated response window (left, center, or right) to receive a liquid reward. Once the rat reached at least 90 trials within 45 minutes for two consecutive days, it proceeded to the Must Initiate phase of the study (**Figure 3A**), in which the light above the liquid reward dispenser was illuminated and the rat was required to insert its nose into the feeder to initiate a trial. This resulted in the feeder light turning off, a tone, and one of the three windows on the touchscreen being illuminated. If the rat touched its nose to the illuminated response window on the screen a reward was delivered. Once the rat completed more than 80 Must Initiate trials within a 45-minute session, the animal moved to the final stage of shaping (**Figure 3A**). In this final phase, if the rat touched the illuminated window, the feeder light illuminated, and a liquid reward was delivered. Conversely, if the rat touched a non-illuminated window, no reward was made available, a tone played to indicate the error, and the house light was illuminated in the box for a four-second timeout. After the timeout and the five-second inter-trial interval, the rat was then able to initiate the next trial. Once the rat achieved over 80% correct of total trials on Punish Incorrect for two consecutive days, it moved on to the procedural training for the OSCS task.

**Figure 3.**
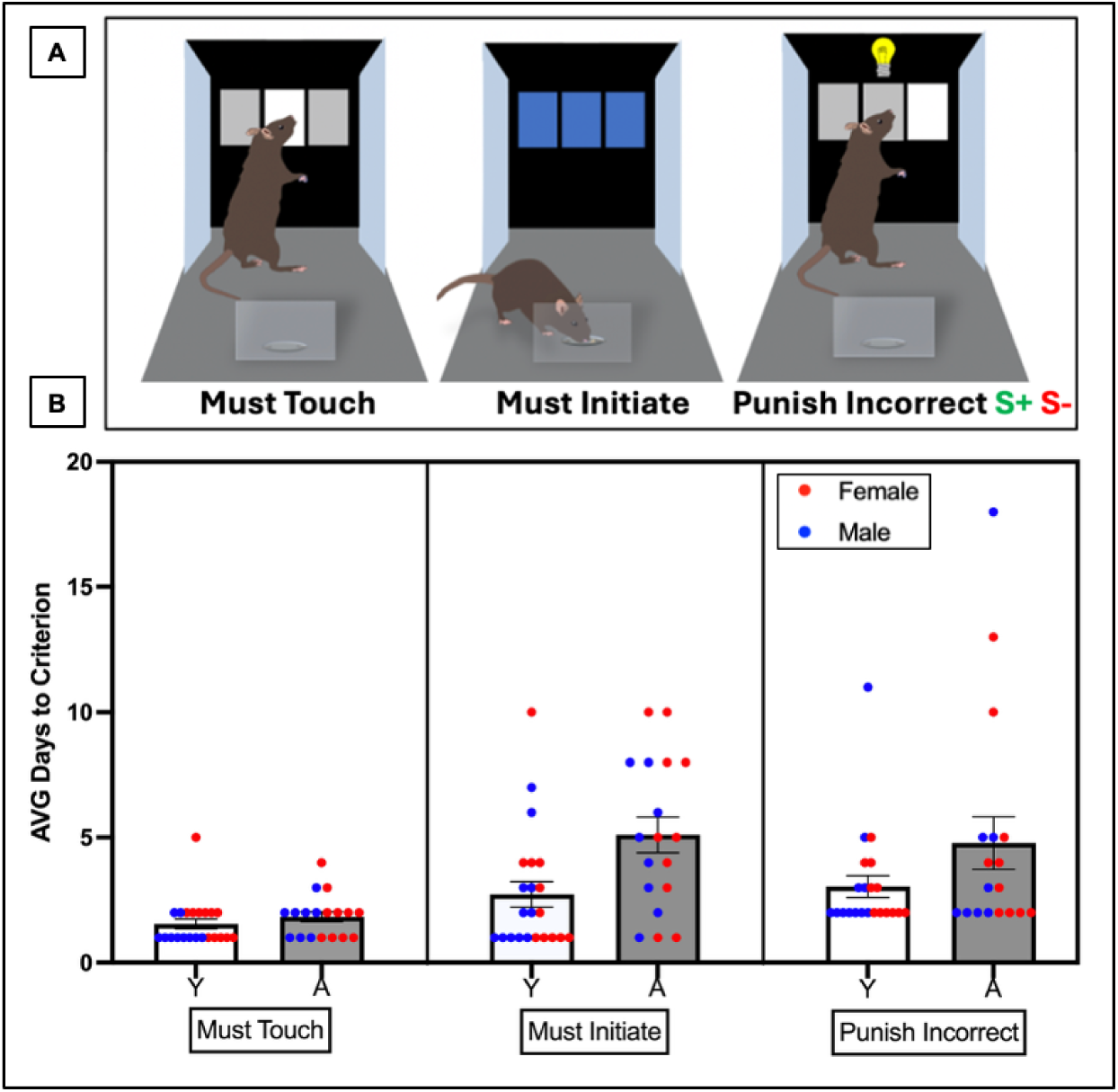
Must Initiate, Must Touch, and Punish Incorrect Shaping. (A) Representative images illustrating three advanced shaping phases: Must Initiate, Must Touch, and Punish Incorrect. In the Must Initiate phase, the rat learns to begin each trial by placing its head into the illuminated food reward dispenser. The Must Touch phase requires the rat to respond specifically to a single illuminated response window to receive a food reward. In the Punish Incorrect phase, incorrect responses (touching a non-illuminated or incorrect window) result in a brief timeout accompanied by the house light turning on to cue that an incorrect choice was made. (B) Average number of days required to reach criterion for each shaping phase in young (Y) and aged (A) rats. Data are presented as mean ± SEM with individual data points overlaid and color-coded by sex.

### Object-Cued Spatial Choice (OCSC) Task

Prior to testing on the morphed OCSCT all rats were trained on the OCSC task (Ahn and Lee, 2015) with two different easily distinguishable black and white image sets **(Figure 4A).** This was done to measure whether or not there were potential age differences in the sensorimotor requirements needed to perform touchscreen-based visual discrimination tasks and to familiarize rats with the basic procedural requirements of the morph-based OCSC task. In the OCSC task rats initially learned to respond to a specific image stimulus that was presented in the center response window via Cued training **(Figure 4B-left)**. The rat was required to touch the image, and then a cue illuminated in one of the response windows (either left or right depending on the image) to indicate the correct choice between the left versus right window. One image of a testing pair was always associated with a left cue while the other image was always associated with a right cue. Touching the cue resulted in the delivery of the reward. If the rat failed to select the cue, the house light was illuminated, and a timeout of four seconds began. Once the rat achieved over 80% correct responses for each trial type (left cue and right cue) for at least two consecutive days, it moved on to the next stage of OCSC training, Non-cued training **(Figure 4B-right)**, in which the side associated with the correct choice for a given image was not indicated and cues appeared on both the left and right sides to designate the response requirement without indicating the correct choice.

**Figure 4.**
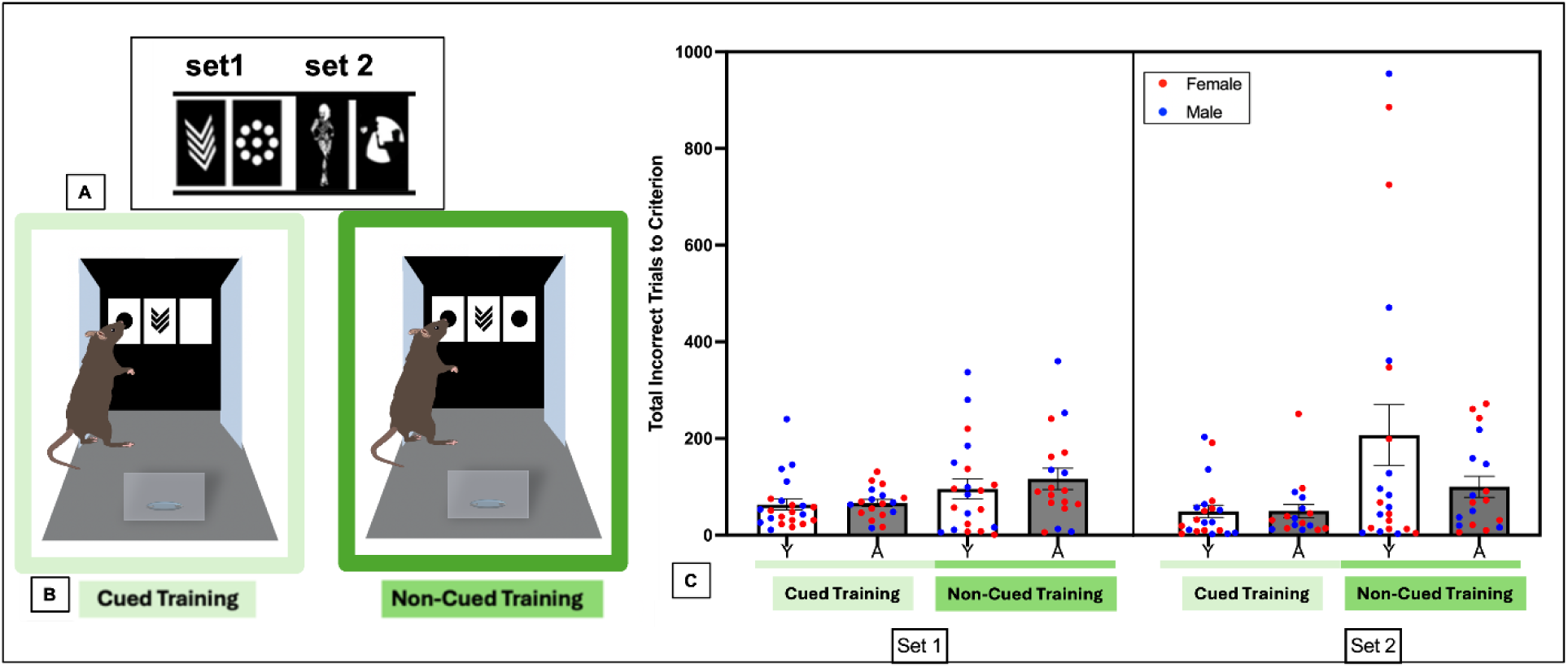
Procedural Training phase of the Object-Cued Spatial Choice (OCSC) task. (A) Stimulus sets used during the OCSC training phase. (B) Representative images illustrating the two training conditions: Cued (left) and Non-Cued (right) training. (C) Total number of incorrect trials required to reach criterion for each training condition (Cued vs. Non-Cued) across stimulus sets (1-2) in young and aged rats. Data are presented as mean ± SEM with individual data points overlaid and color-coded by sex.

Selection of the correct position resulted in the availability of a liquid reward, while selection of the incorrect screen position resulted in a timeout for four seconds with the house light illumination. Once the rat achieved over 80% correct responses for each trial type (chevron and nine) for at least two consecutive days, it moved on to the next stimulus set (Ru/Klaus **Figure 4A**-set 2). To assess potential age differences in procedural training on the OCSC task, a repeated-measures ANOVA was conducted on the number of incorrect trials with Set (1 versus 2) and Cued condition as within-subject factors and Sex, and Age as between-subject factors. There was a significant main effect of Cued condition (F[1,33] = 12.983, p< 0.001, η²₉ = 0.28), indicating that significantly fewer errors were made when the correct OCSC response was cued compared to non-cued. No significant effects were observed for Sex or Age, nor interactions involving these factors (all p> 0.05).

During testing on the Morph OSCST, rats were initially trained to discriminate between two dissimilar images (i.e., bulldog-owl; **Figure 5A/B**) with the same procedural training used for the Set 1 and 2 images with the left versus right image association first being cued and then non-cued training. As in procedural shaping, the image-side association was counterbalanced across rats. Rats were required to reach 80% or more correct responses for each image before proceeding onto the last phase of the study in which performance was tested by morphing the two stimulus images to varying levels of similarity. During the final testing stage on the Morph OCSC task, the stimuli presented included the pre-trained bulldog and owl images and six additional lure images (**Figure 5**). These lure images were generated with Morpheus Photo Mopher software (Howell, MI) to have overlapping features with the original images that varied from 12% to 88% pixel overlap with the original images. The image with the greatest feature overlap indicated the side that was the correct choice for receiving a reward. For example, if an image had 12% overlap with the bulldog and 88% overlap with the owl, the correct choice would be to touch the response window associated with the owl. Conversely, if an image had 64% overlap with the bulldog and 36% overlap with the owl, the correct choice would be to select the response window associated with the bulldog. In each morph session of the OCSC task, there were eight possible trial types composed of the following feature overlap conditions: 1) 100% owl/0% bulldog, 2) 88% owl/12% bulldog, 3) 75% owl/25% bulldog, 4) 64% owl/36% bulldog, 5) 36% owl/64% bulldog, 6) 25% owl/75% bulldog, 7) 12% owl/88% bulldog, and 8) 100% bulldog/0% owl. Each trial type was presented in a pseudorandom order so that it was not repeated more than two consecutive times, and there were up to 20 trials per overlap condition for a total of 160 trials per morph session, however, rats did not always complete all 160 trials. Therefore, only sessions in which rats completed at least 80 trials (close to 10 trials per condition) were included in the analysis. Note that each feature overlap quantification was used twice (0%, 12%, 25%, 36%), once for similarity to the owl and once for similarity to the bulldog. Thus, performance on trial types with the same overlap values (but two different images) were collapsed for a total of 4 trials. This also allowed for more accurate percent correct per each overlap condition to be calculated by avoiding skewed values to a small number of trials. All rats completed the morph sessions at least three times, and the average performance across these sessions was analyzed for significance.

**Figure 5.**
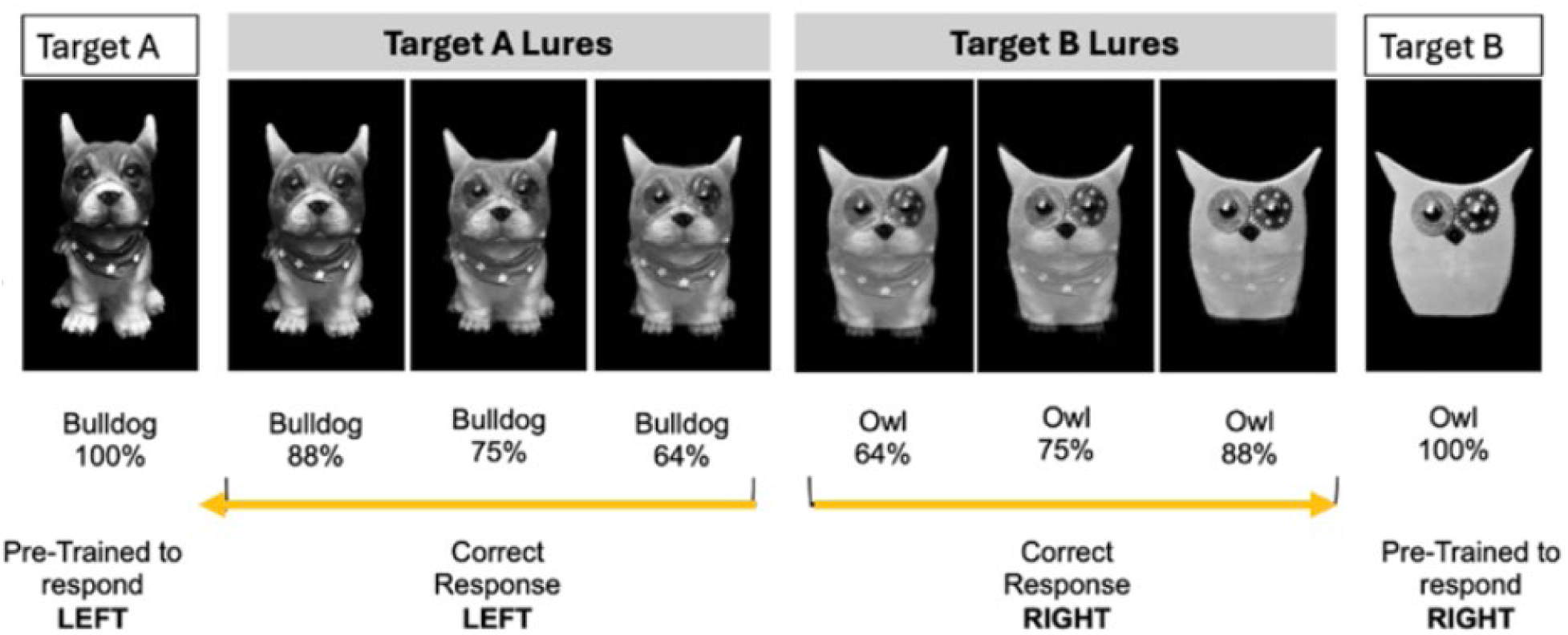
Stimuli used in the Morph OCSC task. The morphed stimulus set used during OCSCT testing. The target images were a Bulldog (Target A) and Owl (Target B). Rats were trained with both Cued and Non-Cued conditions to learn the associated side for each target image. Morphing continuum for each target image generated by blending the Bulldog (Target A) and Owl (Target B) images at varying similarity levels from 100% to 64%, including: 1) Trial type 1 - 100% (0% feature overlap), 2) Trial type 2 - 88% (12% feature overlap), 3) Trial type 3 - 75% (25% feature overlap), and 4) Trial type 4 - 64% (36% feature overlap).

### Cellular Compartment Analysis of Temporal Activity with In situ hybridization (CatFISH) for the Immediate-early gene Arc

To determine is the perirhinal cortex was similarly engaged across feature overlap conditions in young and aged rats, a subset of young (4-month, n = 5♀/4♂) and aged (21-month, n = 3♀/2♂) rats were tested the OCSC task to compare perirhinal cortical principal cells activity during high overlap versus low overlap conditions. Behavioral testing took place in the touchscreen chamber as previously described, however, to isolate activity associated with low versus high feature overlap each animal performed a single stimulus overlap condition for 5 minutes, followed by a 20 min rest in the home cage, and then a final 5 minute epoch with the other stimulus overlap condition prior to immediate sacrifice. Stimulus similarity served as the primary manipulation of task difficulty. In the low overlap condition (Low OCSCT), images were visually distinct with 0% feature overlap (**Figure 6A-left**), whereas in the high overlap condition (High OCSCT), images shared 25% overlapping features (**Figure 6A-right**). Each rat completed two 5-minute OCSC task sessions within a single testing day. The first session was followed by a 20-minute inter-session interval in the home cage, after which animals completed a second 5-minute session with the task difficulty reversed (Low → High or High → Low). This design enabled the detection of RNA for the immediate-early gene *Arc* to be used to infer neuronal activity associated with low versus high stimulus overlap, consistent with the Cellular Compartment Analysis of Temporal Activity with In situ hybridization (CatFISH) (Guzowski et al., 1999).

**Figure 6.**
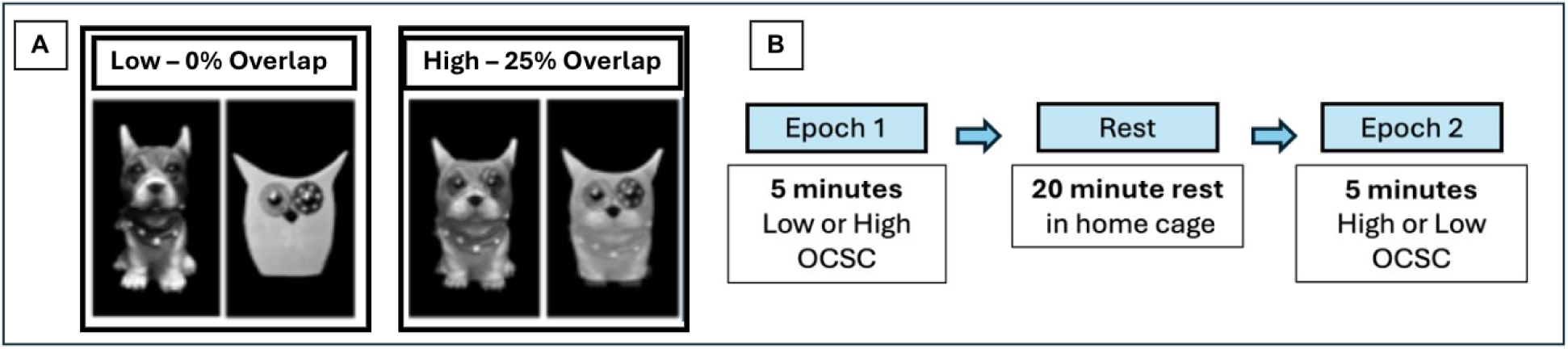
Experimental design and stimulus conditions for the *Arc* CatFISH modified OCSC task. (A) Example image pairs used in Modified OCSCT. Low condition stimuli were visually distinct with 0% feature overlap, whereas High condition stimuli shared 25% overlapping features. (B) Schematic of the within-subject testing design. Rats completed two 5-minute OSCT sessions separated by a 20-minute home cage interval.

Immediately following the second testing session, animals were deeply anesthetized in a glass jar containing isoflurane-saturated cotton separated from the animal by a wire mesh shield. After loss of righting reflex animals were sacrificed via rapid decapitation. For positive control, 4 rats received maximal electroconvulsive shock (MECS) for 5 or 30 minutes prior to sacrifice. Tissue was extracted, hemi-sected, and flash-frozen in 2-methylbutane chilled in a bath of dry ice with 100% ethanol (∼ −70 C). Tissue was stored at −80◦C until cryosectioning. Tissue blocks were made that contained brains from each experimental group to control for slight variations in tissue labeling across slides. These tissue blocks where sectioned at 20 µm on a cryostat (Microm HM550) and thaw mounted on Superfrost Plus slides (Fisher Scientific). Sliced tissue was stored at −80◦C until in situ hybridization.

To measure activity-related gene expression, in situ hybridization was performed to label *Arc* mRNA for catFISH as previously described (Guzowski et al., 1999). A probe was made from the *Arc* gene sequence and labeled with a chemical tag (digoxigenin, DIG, REF #: 11277073910, Roche Applied Science) so it could later be visualized. This probe was applied to brain sections and allowed to bind overnight to *Arc* mRNA present in the cells. After washing, the bound probe was detected with anti-DIG (Catalog #700772, Invitrogen). The *Arc* signal was revealed with a cyanine-3 detection kit (TSA Cyanine 3 Tyramide; PerkinElmer, REF#: 10197077), and nuclei were counterstained with DAPI (Invitrogen) to visualize the total number of cells. This approach allowed us to identify neurons that had recently been active (red) within the broader population of cells (blue).

After staining, z-stacks at 1 μm increments were collected by fluorescence microscopy on a Keyence BZX-810 digital microscope (Keyence Corporation of America, Itasca, IL). Four images were taken from the deep and superficial layers of the perirhinal cortex. Experimenters quantified *Arc* expression in neurons using ImageJ software with a custom written plugin for identifying and classifying cells (available upon request). Only cells that were completely within the frame of the image and within the median 20% of the optical planes were included. The total number of cells were identified before the *Arc* color channel was visible so that signal in these channels did not bias initial segmentation results. Cells were classified as positive for nuclear *Arc* (indicating neuron spiking 1-5 min before sacrifice), cytoplasmic *Arc* (indicating neuron spiking 25-20 min before sacrifice), both nuclear and cytoplasmic *Arc* (activity during both epochs), or negative for *Arc* (no activity in the 30 minutes prior to sacrifice). Neurons were classified based on *Arc* signal localization as follows: 1) Cells were considered nuclear (Foci+) if one or two bright, punctate signals were present within the nucleus and visible across at least three consecutive optical planes. Cells were classified as cytoplasmic (Cyto+) if *Arc* signal appeared as a speckled distribution surrounding ≥50% of the nucleus across multiple planes, without nuclear foci present. Cells meeting both nuclear and cytoplasmic criteria were designated Both+, while neurons not meeting any of these criteria were classified as negative for *Arc* expression. *Arc* in the cytoplasm indicates that the cell fired during the first segment of OCSC testing (epoch 1, Figure 4B), as it takes approximately 20–30 min for the mRNA to translocate to the cytoplasm. Arc foci in the nucleus indicate that the cell fired during the second segment of OCSC testing (epoch 2, Figure 4B). Animals were excluded from analyses if tissue damage prevented reliable quantification or if fewer than two valid counts were obtained for any perirhinal cortex subregion (area 35 or 36; superficial or deep layers).

### Statistical analyses

All statistical analyses were conducted in *RStudio* (R version 2022.12.0; R Core Team) using relevant packages for mixed modeling and analysis of variance. Data were screened for normality, homogeneity of variance, and outliers prior to significance testing. To assess group differences in behavioral performance, mixed-factorial analyses of variance (ANOVAs) were conducted with age (young vs. aged) and sex (male vs. female) included as between-subjects factors, and stimulus overlap as the within-subject repeated measures. When assumptions of sphericity were violated, Greenhouse Geisser corrections were applied. Linear mixed-effects models (LMMs) were additionally used to account for repeated measures and subject-level variability, with subject included as a random effect. Fixed effects included age, sex, and task-related factors, as appropriate. Model fit was evaluated using marginal and conditional R² values, reflecting variance explained by fixed effects alone and by the full model, respectively. To examine relationships between neural measures and behavior, general linear models were conducted with performance metrics as the dependent variable and *Arc* expression levels or similarity score (SI) measures as predictors. age and sex were included as covariates where appropriate. Significant main effects and interactions were followed by post hoc comparisons using estimated marginal means with Tukey-adjusted p-values. Effect sizes are reported as eta squared (η²) for ANOVAs and corresponding measures for mixed models where applicable. Statistical significance was set at α = 0.05. All data are presented as mean ± standard error of the mean (SEM).

## RESULTS

### Morph Object-Cued Spatial Choice (OCSC) Task Performance

To assess for potential age and sex effects on acquiring the image-side association, a mixed ANOVA examining incorrect trials for the OCSC set (Bulldog-Owl, **Figure 7**) was employed. Surprisingly, aged rats made significantly fewer errors overall than did the young animals (F[1,36] = 4.71, p = 0.0163, η²₉ = 0.12). There was also a significant main effect of cue condition (cued or non-cued) (F[1,36] = 9.92, p = 0.0047, η²₉ = 0.22). Unlike with the training images, rats made more errors before reaching criterion for the cued version compared to the un-cued version. This was more evident in the young rats as indicated by the significant age x cue interaction (F(1,36) = 5.80, p = 0.021, η²₉ = 0.14). This observation suggests that all rats were able to generalize the OCSC task rule across stimulus sets.

**Figure 7.**
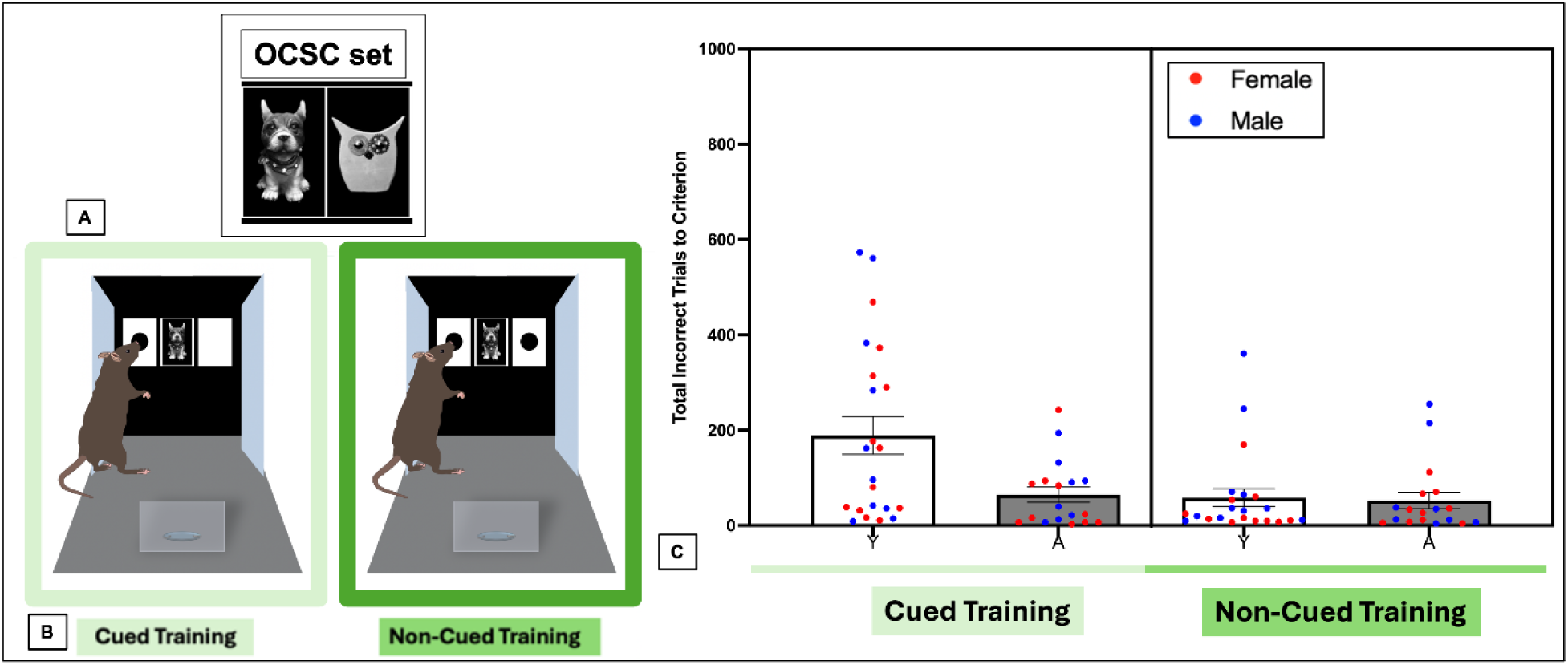
Stimulus Training for the OCSC Task. (A) Owl-bulldog stimulus set used during the OCSC testing phase (left-Target A [Bulldog], right-Target B [Owl]). (B) Representative images illustrating the two training conditions for OCSC set: Cued and Non-Cued training. (C) Total number of incorrect trials required to reach criterion for each training condition (Cued vs. Non-Cued) in young and aged rats. Data are presented as mean ± SEM with individual data points overlaid and color-coded by sex.

**Figure 8.**
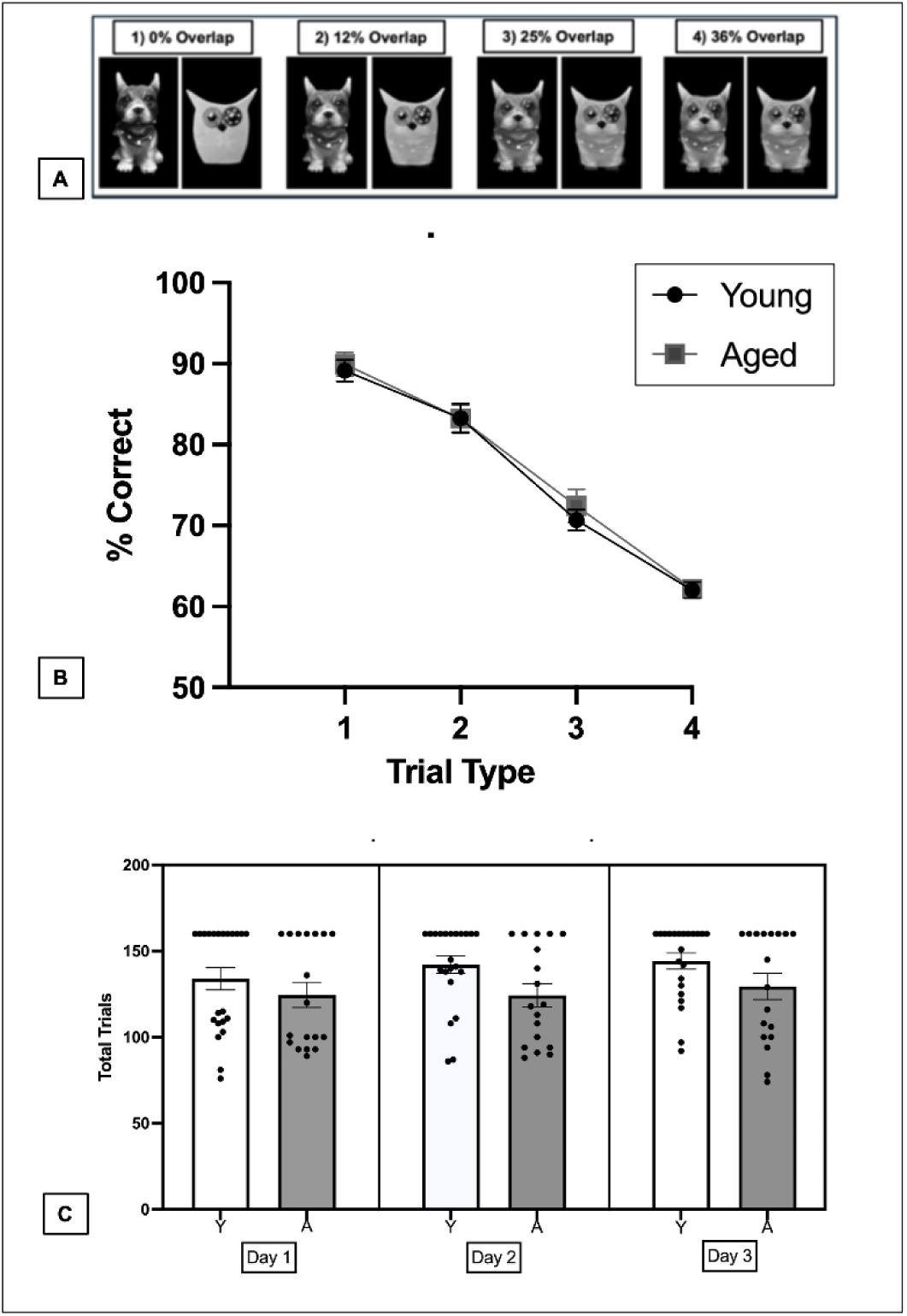
Average performance on the Morph OCSC task. A) (Example images used for each trial type: Trial Type 1 consists of the original target images, while Trial Types 2-4 feature progressively morphed images with increasing similarity to the target. Data are presented as mean ± SEM. B)) The average percent correct for each trial type (T1-T4) across three days of testing. Performance decreases as image similarity increases, with Trial Type 1 (T1, target images) showing the highest accuracy and Trial Type 4 (T4, most similar morphed images) showing the lowest accuracy.(C) The average total trials completed each day of testing between young and aged rats C) Please add the supplement figure of trials completed across day to this panel.

To examine how morphing the Owl and Bulldog stimuli impacted OCSC task performance, a repeated measures ANOVA was conducted on average percent correct across the three morph testing days to examine the effects of Trial Type (T1: 0% overlap, T2: 12% overlap, T3: 25% overlap, and T4: 36% overlap), age, and sex. There was a significant main effect of trial type (F[3,108] = 237.182, p < 0.001, η²₉ = 0.82), indicating that performance significantly decreased as a function of stimulus similarity. No significant main effects or interactions were observed for age, sex, or group (p > 0.05 for all comparisons). Post hoc comparisons (Tukey-adjusted) revealed that performance was highest for T1 (low similarity) and progressively declined with increasing similarity (T2-T4; p < 0.001 for all comparisons), indicating that discrimination performance decreased as similarity increased independent of age, or sex.

One possibility for the lack of an observed age effect is that the aged rats improved across testing days. To evaluate whether practice effects mitigated potential age differences, a repeated measures ANOVA was conducted to analyze performance (%correct) across trial types (T1-T4; within subjects factor) and testing days (D1-D3; within subjects factor), with sex and age group as between subjects factors. Although the main effect of trial type on performance persisted, (F[3,108] = 161.58, p<0.001, η²₉ = 0.82). There was no significant main effect of day (F[2,72] = 1.929, p= 0.15), suggesting consistent performance across days.

The lack of an age effect could also potentially be related to differences in the numbers of completed trials across testing. To examine this possibility, a repeated measures ANOVA was conducted to compare differences in the number of completed trials across testing days (Day 1-3) and to assess effects of sex, and age (**Fig-8C**). There was no significant main effect of day, (F[2,72] = 1.689, p = 0.19), sex (F[1,36] = 0.290, p = 0.59), or age (F[1,28] = 2.82, p = 0.10) on the numbers of trials completed. Additionally, there were no significant interactions between sex, age or day (p > 0.05 for all comparisons).

Previous studies have reported that when faced with a two-choice discrimination with similar 3-dimensional objects, aged rats are more likely to have a bias to select an object on one side regardless of the stimulus (Johnson et al., 2016; Johnson et al., 2017; Colon-Perez et al., 2019). In the OCSC task, this type of response bias would be reflected as a consistent preference for one side of the touchscreen, which can be quantified with a side bias index calculated as: [abs. value (total left touches – total right touches) ÷ total touches]. A score of 0 indicates no side bias, while a score of 1 reflects the maximal side bias (that is, selecting the same side of the screen for every trial). **Figure 9** depicts the mean side bias scores across trial types. A repeated-measures ANOVA revealed no significant main effect of trial type on side bias (F[2.31, 57.77] = 2.37, p = 0.09). There was also no significant main effect of age (F[1, 25] = 1.88, p = 0.18) or sex (F[1, 25] = 2.39, p = 0.13). Additionally, no significant interactions were found between trial type and age (F[2.31, 57.77] = 0.12, p = 0.91), trial type and sex (F[2.31, 57.77] = 2.51, p = 0.82), or age and sex (F[1, 25] = 2.94, p = 0.10), nor was the three-way interaction significant (F[2.31, 57.77] = 0.55, p = 0.60). Together, these results indicate that side bias did not significantly differ across trial types, age groups, or sexes, even under high similarity conditions.

**Figure 9.**
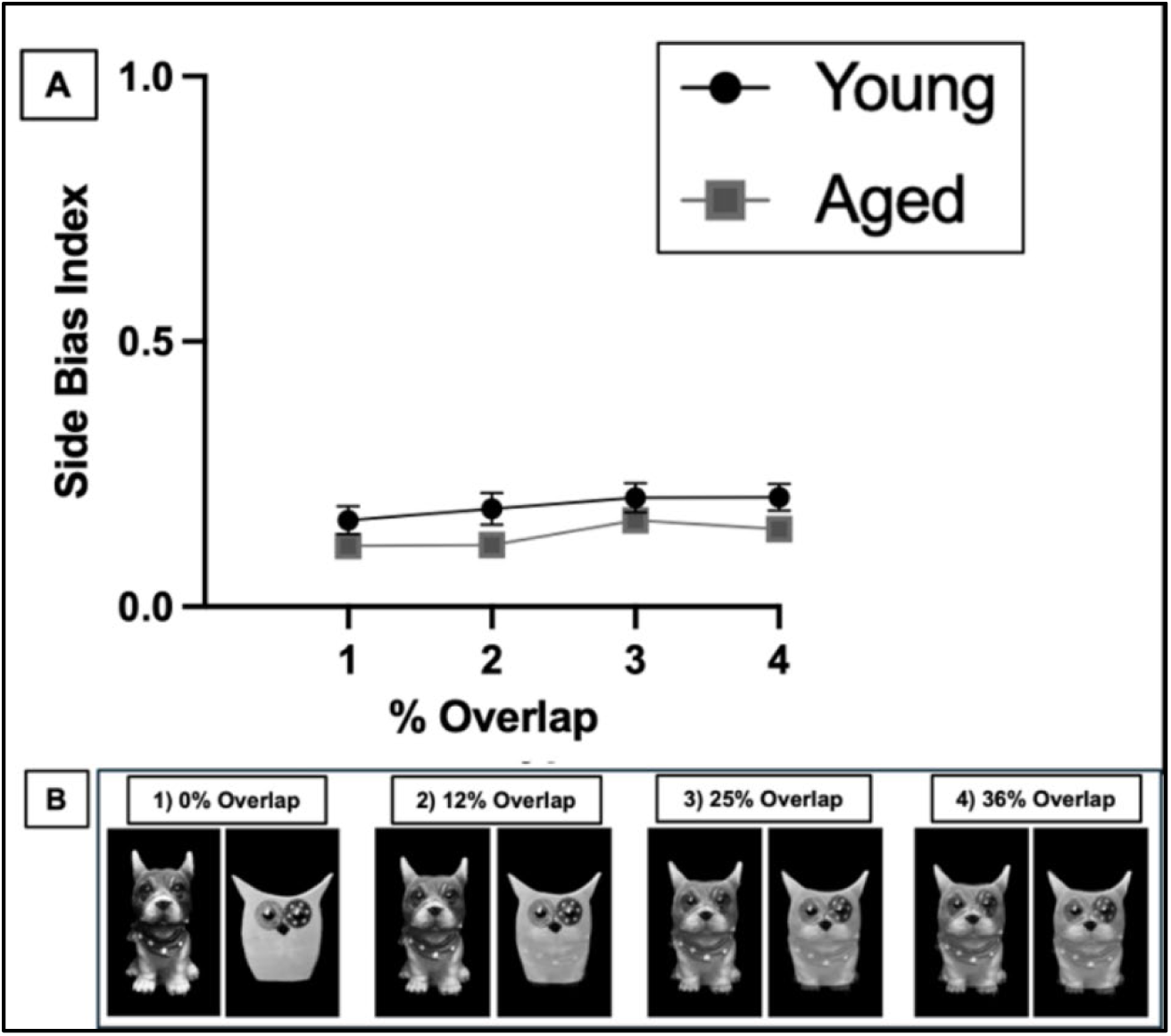
Side bias index on the Morph Object-Cued Spatial Choice Testing task. (A) The mean side bias index for each trial type (T1-T4) across three days of testing. Values closer to 0 indicate less bias, while values closer to 1 indicate a stronger side preference. (B) Example images used for each trial type: Trial Type 1 consists of the original target images, while Trial Types 2-4 feature progressively morphed images with increasing similarity to the target. Data are presented as mean ± SEM.

### Perirhinal Cortical Arc Expression During OCSC Task Low and High Overlap Conditions

To evaluate whether the perirhinal cortex was similarly engaged during low and high overlap conditions in the OCSC task, a modified version of the task was implemented with a subset of rats in which rats were tested on the OCSC task with no feature overlap for all trials during one 5-min epoch of behavior and in a different 5-min epoch, the morphed images with 25% feature overlap were used for all trials (images shown in **Figure 10A**). The different overlap epochs were separated by 20 min with the order counterbalanced so that the subcellular location of the *Arc* mRNA could be used to infer ensemble activity during the two different overlap conditions using the Cellular Compartment Analysis of Temporal Activity by Fluorescence *in situ* Hybridization (CatFISH) (Guzowski et al., 1999). The perirhinal cortex was selected for this analysis because it has been previously reported to be necessary for performance on the OCSC task (Ahn and Lee, 2015), and age-related deficits in stimulus discrimination and object recognition have been linked to dysfunction in this structure (Burke et al., 2012; Burke and Barnes, 2014; Maurer et al., 2017; Burke et al., 2018b; Hernandez et al., 2018).

**Figure 10.**
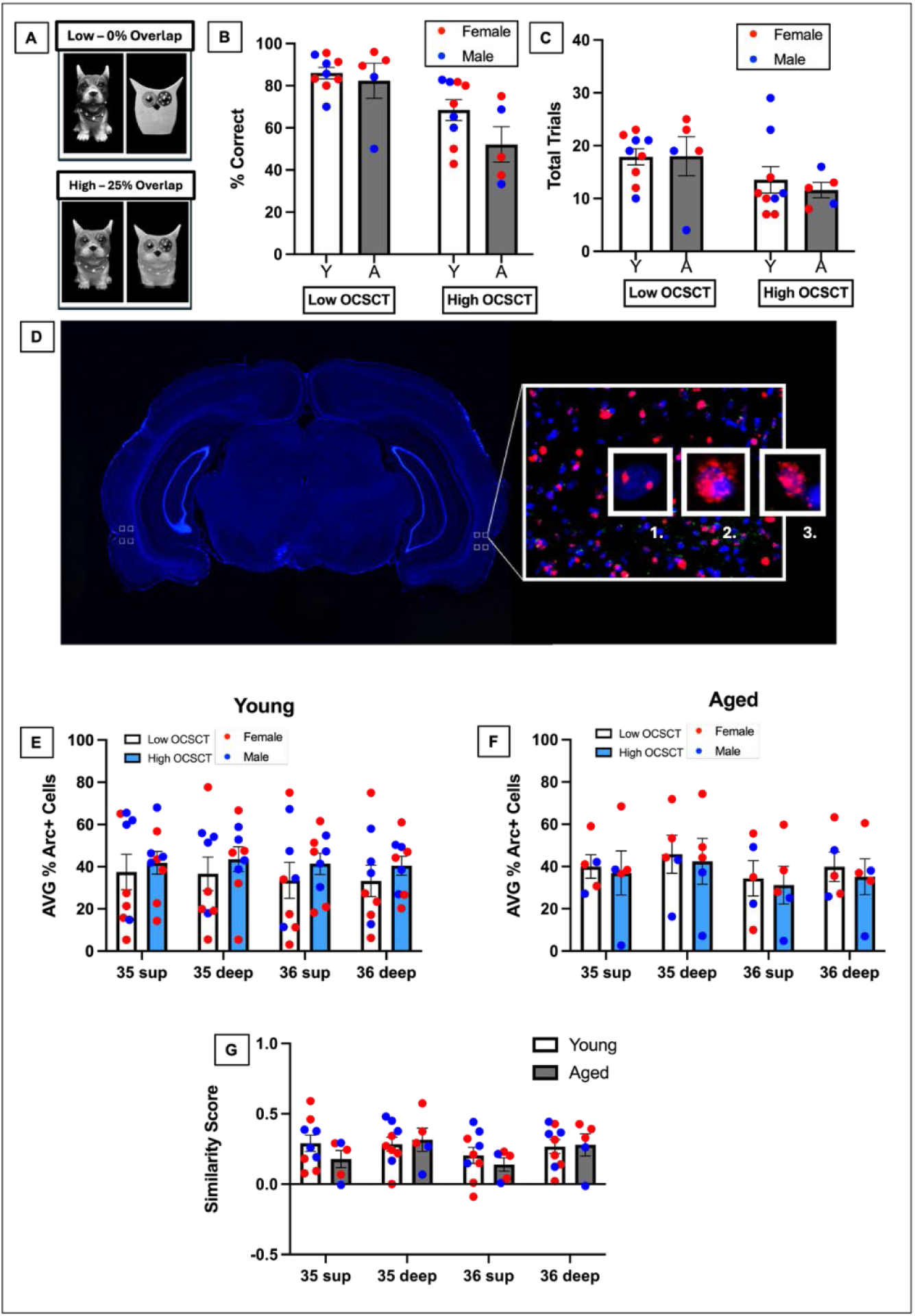
Behavioral performance and Arc expression in the Modified OCSC Task. (A) Example stimulus pairs used to manipulate task difficulty. Low OCSC task stimuli have 0% feature overlap, whereas High OCSCT stimuli shared 25% overlapping features. (B) Average percent correct on Low and High OCSCT in young and aged rats. (C) Average total trials completed by young and aged rats across overlap conditions. (D) Representative coronal brain section indicating the areas of perirhinal cortex (PRC) that were imaged for *Arc* expression analysis, including superficial and deep layers of areas 35 and 36. (D1–D3) Representative fluorescence images illustrating *Arc* expression patterns (red) for 1) foci only (activity 2-5 min before sacrifice), 2) cytoplasm only (activity 25-30 min before sacrifice), and 3) both foci and cytoplasmic *Arc* (activity during both epochs). Blue is DAPI-stained nuclei. (E) *Arc*-positive cells in superficial and deep layers of perirhinal areas 35 and 36 in young rats. (F) *Arc* expression in aged rats. (G) Similarity Score in superficial and deep layers of perirhinal area 35 and 36 in young and aged rats. Bars represent group means ± SEM, with individual rats shown as points color-coded by sex.

Behavioral performance was analyzed using a mixed-design repeated measures ANOVA, with Stimulus Overlap as the within subjects factor (0% versus 25% overlap) and age and sex as the between subjects factors (**Figure 10B**). Analysis of discrimination accuracy revealed a significant main effect of stimulus overlap, indicating that performance was higher in the 0% overlap epoch (Figure 10B; F[1,10] = 26.74, p < 0.001, η²₉ = 0.73). However, there were no significant main effects of age (F[1,10] = 2.74, p = 0.129, η²₉ = 0.22) or Sex (F[1,10] = 0.60, p = 0.458, η²₉ = 0.06). Furthermore, the age × sex (F[1,10] = 1.42, p = 0.261, η²₉ = 0.12) and sex × stimulus overlap (F[1,10] = 3.42, p = 0.094, η²₉ = 0.25) interactions were also not significant. The observation of an effect of stimulus overlap with no age effect indicates that performance on the modified OCSC task was similar to the full morph OCSC task used to probe behavioral effects.

A mixed-design ANOVA was conducted on total trials completed during the 5 min epoch to assess whether the factors of Stimulus overlap, age, and Sex influenced the completion rate (**Figure 10C**). There were no significant main effects of age (F[1,10] = 0.64, p = 0.442, η²₉ = 0.06) or Sex (F[1,10] = 0.02, p = 0.891, η²₉ = 0.00) or Stimulus Overlap (F(1,10) = 3.36, p = 0.097, η²₉ = 0.25). Moreover, none of the interaction effects reached statistical significance (p >0.05 for all comparisons). Together these observations indicate that the number of trials completed is not likely to contribute to potential differences in perirhinal cortical *Arc* expression.

The proportion of perirhinal cortical neurons expressing *Arc* in association with OCSC task performance was analyzed using a linear mixed-effects model (LMMs) with Overlap condition (Low vs High OCSC), Region (area 35 vs 36), and Layer (superficial vs deep) as within-subjects factors and age and Sex treated as between-subject factors. *Arc* expression did not differ across Stimulus Overlap conditions (F[1,70] = 0.0190, p = 0.891), and there were no significant main effects of age (F[1,10] = 0.113, p = 0.744), Sex (F[1,10] = 0.223, p = 0.646, Region (F[1,70] = 1.9846, p = 0.167) or Layer (F[1,70] = 0.497, p = 0.483). Group averages and individual animal distributions are shown in **Figure 10E** (Young rats) and **Figure 10F** (Aged rats).

To determine whether neural activation predicted behavioral performance, we examined the relationship between *Arc* expression and behavioral performance using a linear mixed-effects model with percent correct as the dependent variable and *Arc* expression as a fixed effect. This analysis revealed that *Arc* expression was not significantly associated with behavioral performance on the modified OCSC task (β = −0.172, SE = 0.143, t = −1.20, p = 0.234, Marginal R^2^ = 0.029).

A powerful aspect of the *Arc* CatFISH approach is that it allows one to determine the extent to which neuronal populations engaged during 2 different experiences orthogonalize. As population overlap can be affected by differences in overall activity levels, a similarity score can be calculated to control for differences in activity between regions or groups (Vazdarjanova and Guzowski, 2004). The similarity score between the 0% and 25% overlap conditions was therefore calculated as:

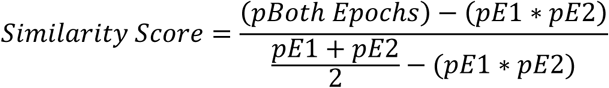

where pBoth Epochs is the proportion of cells active during both epochs (that is, both foci and cytoplasmic *Arc*) and pE1 and pE2 are the proportion of cells active during Epoch 1 (cytoplasm + both) and Epoch 2 (foci + both), respectively. The mean Similarity Score across groups and perirhinal cortical areas is shown in **Figure 10G**. The relationship between Similarity Score and age, sex, regions (area 35 versus 36) and perirhinal cortical layer was analyzed using a mixed factorial ANOVA. No significant main effects of age (F[1,20] = 1.10, p = 0.306, η²₉ = 0.05), Sex (F[1,20] = 0.12, p = 0.735, η²₉ = 0.00), or Region (F[1,20] = 0.68, p = 0.419, η²₉ = 0.03) on the Similarity Score were detected. However, a significant main effect of Layer was observed (F[1,20] = 6.36, p = 0.020, η²₉ = 0.24), with deep layers of the perirhinal cortex having a significantly greater Similarity Score compared to the superficial layers, which has been reported previously (Burke et al., 2005; Takehara-Nishiuchi et al., 2013). To interrogate whether the similarity score was related to the difference in correct trials between the low and high overlap stimulus conditions, a linear mixed-effects models with task performance difference (no overlap %correct – high overlap %correct) as the dependent variable and similarity score as a predictor. Potential moderators (age, sex, cortical region and layer) were then tested in separate models by adding interaction terms with similarity score. Across all models, similarity scores were not significantly associated with performance difference (all p > 0.5). Overall, these results provide no evidence that similarity scores predicted task performance, nor was this relationship is moderated by age, sex, cortical region or layer.

## DISCUSSION

The present study evaluated whether a touchscreen-based, morph Object-Cued Spatial Choice (OCSC) task could capture age-related mnemonic discrimination deficits and whether such performance was associated with perirhinal cortex (PRC) engagement as measured by expression of the immediate-early gene *Arc*. Consistent with prior work, discrimination accuracy declined systematically as feature overlap increased, demonstrating that the task successfully manipulated perceptual similarity. However, contrary to robust findings from rodent and human mnemonic similarity task (MST) paradigms, young and aged rats performed equivalently across all overlap conditions. Moreover, PRC *Arc* expression did not differ as a function of age, stimulus overlap, or sex, revealing a striking dissociation between behavioral sensitivity to stimulus similarity and neural recruitment within the PRC. Together, these findings suggest that while the touchscreen-based morph OCSC task is sensitive to perceptual difficulty, it does not capture the age-related discrimination impairments consistently observed in other MST paradigms, raising questions regarding behavioral strategy, neural engagement, and translational validity.

### Dissociation between behavioral performance and PRC engagement

A notable observation in the current study is the divergence between behavioral performance, which decreased as stimulus overlap increased, and PRC *Arc* expression. Based on prior studies reporting that the PRC is critical for resolving feature ambiguity (Murray and Bussey, 1999; Baxter and Murray, 2001; Murray and Richmond, 2001; Bussey et al., 2002; Bartko et al., 2007b, a), we predicted that increasing feature overlap would influence PRC engagement and that this modulation might differ with age. Instead, although performance declined with increasing overlap, the proportion of PRC neurons that had *Arc* expression remained largely unchanged across conditions and groups. However, it is worth stating that there was only a modest degree of neuronal ensemble overlap in the PRC between the epoch with no overlap and the epoch in which the morphed images had 25% feature overlap as quantified with by the similarity score measure (mean <0.4 for all regions, sexes and age groups) (Guzowski et al., 2004). This degree of orthogonalization in the active PRC neuronal populations across two different experiences is comparable to what is observed with the environment changes (Burke et al., 2012) or the cognitive load is altered (Hernandez et al., 2018) between epochs. This observation may suggest that the small differences in visual input from the different images may have provoked different activation patterns in the PRC, which is consistent with a report that the PRC neurons change in their firing rates in response to a stimulus being morphed into a different image (Ahn and Lee, 2017).

While there was a change in the PRC cells that were activated during the OCSC task during 25% overlap versus no overlap trials, it cannot be ruled out that the PRC was not strongly engaged by the touchscreen OCSC task, even under high-overlap conditions. Unlike 3D object-based MST paradigms, which require animals to explore, integrate multimodal sensory inputs, and resolve complex feature conjunctions, the current task relied on 2D visual stimuli presented on a touchscreen. This reduced sensory richness may have diminished the need for PRC-dependent configural processing (Bussey and Saksida, 2002; Cowell et al., 2006) and/or the association of single visual features into a complex percept (Zhang and Zhang, 2026), thereby limiting recruitment of PRC neuronal ensembles. In this framework, the observed behavioral decline with increased similarity may reflect simple perceptual difficulty rather than discrimination of complex multi-feature stimulus configurations that require PRC engagement.

An alternative, but related, interpretation is that PRC involvement in this task is present but not dynamically modulated by increasing mnemonic demand in the same way as in traditional MST paradigms. The absence of a main effect of feature overlap on *Arc* expression suggests that PRC activation may not scale with stimulus similarity under these conditions. However, the presence of a trend-level interaction between age and overlap condition, along with a sex × overlap interaction, raises the possibility that subtle, demand-dependent modulation of PRC activity may exist but is not robustly captured in the current dataset. Indeed, visual inspection suggests that young animals may exhibit a modest increase in PRC activation under higher overlap conditions, whereas aged animals fail to show this modulation. If confirmed, this pattern would align with the hypothesis that aging is associated not with reduced baseline activity, but with a diminished capacity to flexibly recruit cortical circuits in response to increasing task demands. This interpretation is consistent with prior work demonstrating age-related alterations in neural ensemble dynamics, including reduced adaptability and altered recruitment patterns across cortical and hippocampal circuits (Burke et al., 2012; Hernandez et al., 2018). Thus, rather than a global reduction in PRC function, aging may impair the ability to appropriately scale neural responses to task difficulty. Importantly, the lack of a statistically significant age effect in *Arc* expression underscores a key limitation: detecting such differences may require tasks that more strongly engage PRC-dependent computations.

### The morph OCSC task did not detect age differences

The absence of an age-related impairment in the touchscreen morph OCSC task stands in contrast to a large body of literature demonstrating robust deficits in mnemonic discrimination across species. In rodents, aged animals show selective impairments when discriminating between similar objects in 3D object-based MST tasks (Burke et al., 2011; Johnson et al., 2017) and in olfactory discrimination paradigms (Yoder et al., 2017). Similarly, in humans, older adults consistently exhibit deficits on the MST when distinguishing similar lures from previously encountered items (Toner et al., 2009; Yassa et al., 2011b; Stark et al., 2013; Stark et al., 2015; Stark and Stark, 2017; Stark et al., 2019). The failure to detect such deficits in the current study therefore warrants careful consideration. Importantly, the lack of age differences was not attributable to practice effects or differences in trial completion, as performance remained stable across testing days and did not differ in the number of trials completed. Together, these observations support the conclusion that the touchscreen morph OCSC task, in its current form, does not capture the age-sensitive mnemonic processes measured by other MST paradigms.

One likely explanation is that the touchscreen morph OCSC task does not sufficiently tax the neural mechanisms underlying mnemonic discrimination. In traditional MST paradigms, animals must integrate distinct features of a complex object to distinguish the target from the lure. It is conceivable that a 2D stimulus is not experienced by rats in a comparable way and rather than using the full configuration of all visual features, the animals simply extract a single feature to solve the task. This may be more akin to a simple sensory threshold detection, which is intact in aged rats (Yoder et al., 2017), or detecting small variations in color or size. The latter single-feature discrimination is intact in older adults (Ryan et al., 2012) and patients with medial temporal lobe lesions that include the PRC (Barense et al., 2007). Thus, both young and aged animals may be able to perform the morph OCSC task using alternative strategies that bypass the need for fine-grained mnemonic discrimination. This idea is consistent with the observation that rats with PRC lesions perform pairwise discrimination of morphed images similarly to control animals (Clark et al., 2011).

### Ethological relevance of pre-clinical models and translational considerations

A central implication of the current findings is the importance of ethological relevance in designing cognitive tasks to have translational validity for human clinical research. While touchscreen-based paradigms offer clear advantages in terms of automation, standardization, and alignment with human testing formats, they may fail to capture critical aspects of rodent cognition that depend on naturalistic sensory processing. Rodents rely heavily on multisensory integration, including tactile input from whiskers, olfactory cues, and depth perception, when interacting with objects in their environment. Traditional 3D MST paradigms leverage these modalities, requiring animals to explore and discriminate objects in a manner that engages PRC-dependent configural processing. In contrast, 2D images presented on a touchscreen provide a limited and potentially unnatural stimulus set for rodents, which may reduce engagement of the neural circuits underlying mnemonic discrimination. This raises an important translational tension: while aligning rodent tasks with human paradigms is desirable, doing so at the expense of ethological validity may compromise the ability to detect meaningful cognitive deficits. The current findings suggest that, at least for mnemonic discrimination, preserving naturalistic stimulus properties may be more critical than matching stimulus format across species. Future efforts to develop translational MST paradigms may benefit from hybrid approaches that combine the advantages of automation with more ecologically relevant stimulus designs.

## Acknowledgements

Funding provided by Evelyn F. McKnight Brain Research Foundation, NIH/NIA research grant 1R01AG049722, and the University of Arizona College of Science and BIO5 Institute

## Use of AI Disclosure Statement

ChatGPT was used to facilitate coding in R and format statistics.

